# Predicting the genetic ancestry of 2.6 million New York City patients using clinical data

**DOI:** 10.1101/768440

**Authors:** Vijendra Ramlall, Kayla M. Quinnies, Rami Vanguri, Tal Lorberbaum, David B. Goldstein, Nicholas P. Tatonetti

## Abstract

Ancestry is an essential covariate in clinical genomics research. When genetic data are available, dimensionality reduction techniques, such as principal components analysis, are used to determine ancestry in complex populations. Unfortunately, these data are not always available in the clinical and research settings. For example, electronic health records (EHRs), which are a rich source of temporal human disease data that could be used to enhance genetic studies, do not directly capture ancestry. Here, we present a novel algorithm for predicting genetic ancestry using only variables that are routinely captured in EHRs, such as self-reported race and ethnicity, and condition billing codes. Using patients that have both genetic and clinical information at Columbia University/ New York-Presbyterian Irving Medical Center, we developed a pipeline that uses only clinical data to predict the genetic ancestry of all patients of which more than 80% identify as other or unknown. Our ancestry estimates can be used in observational studies of disease inheritance, to guide genetic cohort studies, or to explore health disparities in clinical care and outcomes.

## 1 Introduction

Genetic ancestry is a significant factor in disease pathogenecity and a key pillar in precision medicine^1^. Individuals have genomic variants unique to themselves, their close relatives, and their ancestral populations. Measuring and characterizing the relationship between these variants and clinical factors is now routine when genetic data are available. With existing genetic data, ancestry is estimated using principal components analysis (PCA). PCA reduces dimensionality while maintaining variability, making shared ancestry in complex populations identifiable^2^. Despite the rapid decline in genotyping/sequencing costs^3^, genetic data are not available for the majority of patients in databases of clinical records. This issue is compounded by the time-consuming nature of genotyping/sequencing patients en masse and complicated by issues of consent for re-use. The lack of ancestry data can adversely affect analyses of clinical records. Many diseases and clinical outcomes have unidentified subgroups and variants unique to these populations that can go undiscovered if not properly stratified by ancestry^4^.

An interim goal is to use data that is routinely collected by hospitals to systematically predict the genetic ancestry of patients across an electronic health record (EHR), which routinely collect conditions and drug exposures variables. At Columbia University/ New York-Presbyterian Hospital Irving Medical Center (CU/NYPIMC) we have access to medical records from more than six million patients in a clinical data warehouse (CDW). The diversity present in our CDW, in terms of race, ethnicity, and socioeconomic status, as well as the depth of clinical data collected (over 20 years), makes it an ideal resource for phenotype selection and subgroup identification^5^(Table 1). We hypothesized that these clinical data could also be used to accurately predict genetic ancestry. Essentially, if two clinical concepts are correlated in the EHR and one is known to have a genetic association, then those genetics may also apply to the second concept^6^. Some variables collected in the EHR have clear relationships to genetics, such as self-reported ethnicity and familial relationships^7^, while others are more obscure, such as hypersensitivity due to exposure to the preventative HIV drug abacavir is associated to the genetic variant HLA-B*5701, which is in turn associated with ancestry^8,9^. To evaluate this, we collected whole exome sequencing (WES) data from the Institute for Genomic Medicine (IGM) at CU/NYPIMC for patients that also had clinical data available. We used data from these patients to determine how clinical data can predict ancestry and to evaluate that relationship.

**Table 1:** Demographics

|  | Training Set | Validation Set | EHR |
| --- | --- | --- | --- |
| **N** | 350 | 376 | 6,377,222 |
| **Age** | 36.4 ± 18.0 | 28.5 ± 20.0 | 54.7 ± 25.8 |
| **Gender** |  |  |  |
| Female | 53.71% | 54.26% | 43.99% |
| Male | 46.29% | 45.74% | 55.82 % |
| Other/Unknown/Other values/Blank | 0% | 0% | 0.29% |
| **Reported Race** |  |  |  |
| American Indian or Alaska Native | 0% | 0.27% | 0.01% |
| Asian | 1.43% | 3.72% | 0.53% |
| Black or African American | 6.00% | 5.85% | 3.45% |
| Other Pacific Islander | 0.57% | 0% | <0.01% |
| White | 46.57% | 42.29% | 10.08% |
| Unknown | 37.43% | 40.43% | 79.39% |
| Other | 8.00% | 7.45% | 2.58% |
| Other values/Blank | 0% | 0% | 3.87% |
| **Reported Ethnicity** |  |  |  |
| Hispanic | 26.00% | 23.40% | 4.31% |
| Not Hispanic | 36.57% | 34.04% | 9.20% |
| Unknown | 37.43% | 42.55% | 86.49% |

While surname and physical address have been previously used to estimate race and ethnicity^10^, the use of clinical variables as genetic surrogates of ancestry is a new approach that has not yet been systematically explored. Broad phenotypes are known to have many specific subgroups^11,12^, yet phenotypes mined from clinical data are often described as a single diagnosis or trait. Clinical resources, such as EHRs, collect detailed descriptions of a patient’s state far beyond a given diagnosis code. Further, these records are connected longitudinally, producing a rich time-line of clinical events for every patient. There is an opportunity to leverage these data to identify patient subgroups.

For this study, we used data from the 1000 Genomes Project (1KGP) to derive a principle component (PC) analysis (PCA) model and then we used this model to project the IGM data into PC space. We then trained a random forest regression (RFR) algorithm to predict the PC values from commonly collected clinical data and demographic data for these patients and evaluated this algorithm on an independent test set of IGM patients. We found that we can significantly predict the first four principal components. We then applied this algorithm to all individuals with sufficient data in the EHR (4.4 million patients). We used these predicted PCs to assign ancestry labels for patients and were able to predict ancestry even when patients did not provide any race or ethnicity information. Our population predictions reflect the census data for New York City. These clinically derived ancestry estimates can be used as genetic proxies in modeling for clinical data analysis or to inform genetic studies.

## 2 Results

### 2.1 Acquiring genetic data and matching to clinical population and defining a PCA model

We obtained 2,504 samples from 1KGP containing 5,685,915 variants and 650 samples from HapMap, which were not already present in 1KGP, with 1,014,313 variants after conversion to the GRCh37/hg19 reference genome build. We obtained 4,132 samples from IGM with 2,293,955 variants. After filtering for a minor allele frequency of at least 10% and a genotyping rate of at least 90%, we identified 5,758 single nucleotide variants (SNV) matching chromosome, position, reference allele, alternate allele in common between the three datasets.

Of the 4,132 sequenced samples available at IGM, 726 patients had structured diagnosis data in the CDW at CU/NYPIMC (a total of 12 self-reported race/ ethnicity codes and 3,437 diagnosis codes after mapping to ICD-10). Samples that were available prior to May 2017 served as the training set for the RFR algorithm (N=350), and the remainder served as the validation set (N=376).

We used the 5,758 common SNVs to train a PCA model on the 1KGP sample and then used the model to project the IGM training set data into PC space (Figure 1). The distribution of the first four PCs of the HapMap and IGM patients in the training set correlated with their reported race (Figure S1).

**Figure 1:**
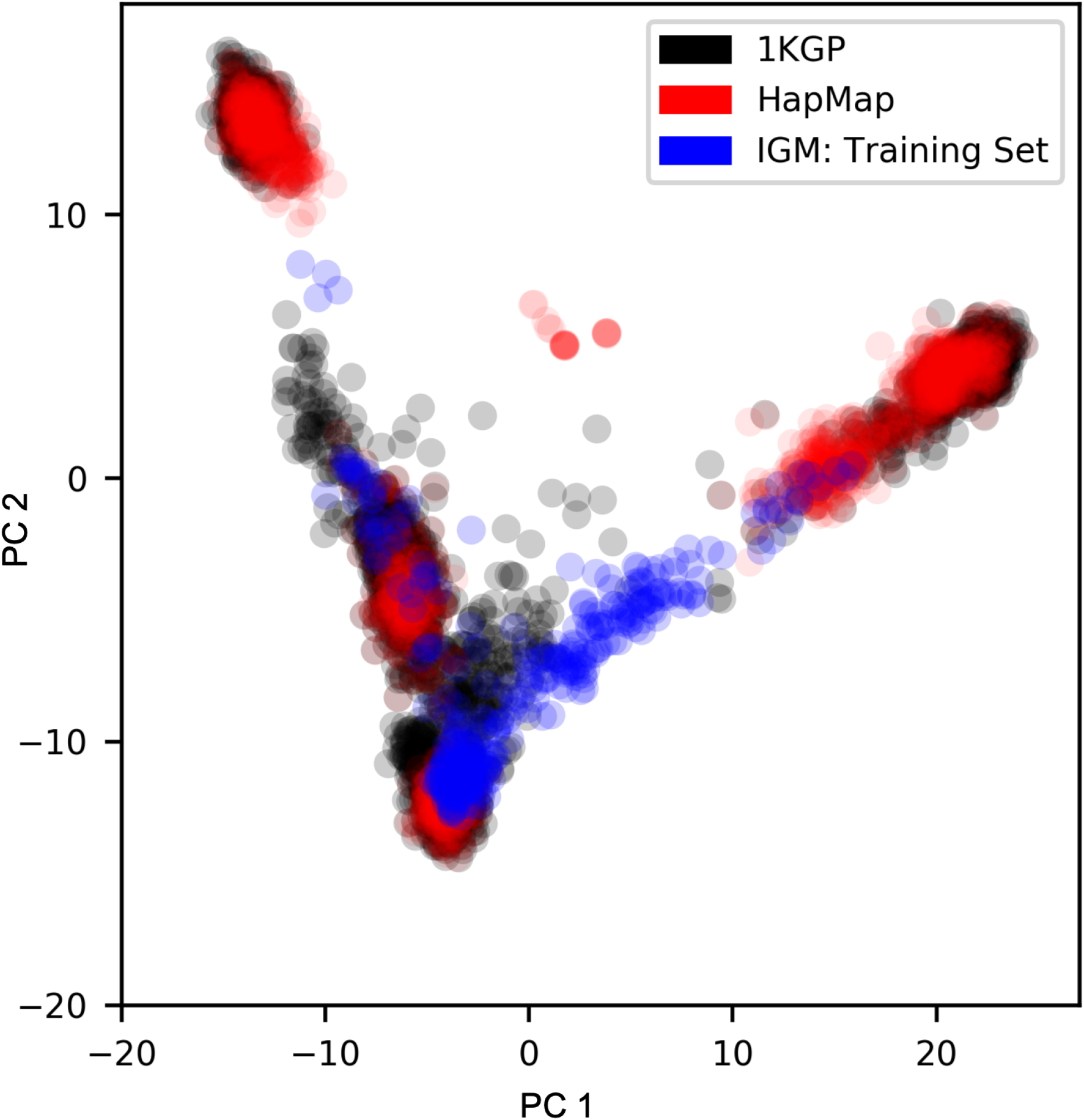
PC 1 vs. PC 2 1KGP, HapMap, and the IGM training set. PC 1 and PC 2 of the PCA of the 1KGP data (black) and PC 1 and PC 2 of the HapMap (red) and the IGM training set (blue) projected using the 1KGP derived PCA model.

### 2.2 Random forest regression algorithm accurately predicts principal components

We used diagnoses and demographic information to predict the genetic principal components. The diagnoses were coded in ICD10 and structured into a hierarchy with four levels (Chapter, Block, Header, Terminal). We trained RFR algorithms on features from each of the four levels of ICD10 hierarchy and found that the first four PCs were predicted from the clinical data regardless of which level’s features were used (Table 2, Figures 2, S2). Of the four levels of the hierarchy, we found that the most abstract level, Chapter, achieved the greatest Pearson correlation, and smallest RMSE, for three of the first fours PCs (Table 2). The PCs predicted by the RFR algorithm trained using the Chapter level features were able to recapitulate the race correlated distribution of the genetics PCs (Figures S3, S4). We therefore used this RFR algorithm in all subsequent analyses. See Table 3 for the 10 most important features of the model.

**Table 2:** Random forest regression model out-of-bag performance at levels of ICD-10 hierarchy

| ICD-10 Level | Pearson correlation | p-value | RMSE |
| --- | --- | --- | --- |
| **Terminal (4172 Features)** |  |  |  |
| PC 1 | 0.6000 | $1.3524 \times 10^{-35}$ | 4.1787 |
| PC 2 | 0.6291 | $5.6620 \times 10^{-40}$ | 3.2371 |
| PC 3 | 0.3079 | $4.0186 \times 10^{-09}$ | 2.1730 |
| PC 4 | 0.3740 | $4.6716 \times 10^{-13}$ | 3.5922 |
| PC 5 | 0.0455 | 0.3963 | 0.7433 |
| PC 6 | -0.0278 | 0.6039 | 1.3280 |
| PC 7 | 0.0754 | 0.1593 | 1.3000 |
| PC 8 | 0.1193 | 0.0257 | 1.1579 |
| PC 9 | -0.0076 | 0.8868 | 1.1982 |
| PC 10 | 0.1507 | 0.0047 | 0.7439 |
| **Header (1816 Features)** |  |  |  |
| PC 1 | 0.5559 | $8.8904 \times 10^{-30}$ | 4.3620 |
| PC 2 | 0.6405 | $8.2876 \times 10^{-42}$ | 3.1776 |
| PC 3 | 0.3000 | $1.0299 \times 10^{-08}$ | 2.2164 |
| PC 4 | 0.3482 | $2.0494 \times 10^{-11}$ | 3.6966 |
| PC 5 | 0.0680 | 0.2043 | 0.7365 |
| PC 6 | -0.0184 | 0.7322 | 1.3232 |
| PC 7 | 0.0688 | 0.1989 | 1.2980 |
| PC 8 | 0.1103 | 0.0392 | 1.1488 |
| PC 9 | 0.0081 | 0.8797 | 1.2128 |
| PC 10 | 0.1557 | 0.0035 | 0.7464 |
| **Block (964 Features)** |  |  |  |
| PC 1 | 0.5573 | $5.9592 \times 10^{-30}$ | 4.3709 |
| PC 2 | 0.6411 | $6.5091 \times 10^{-42}$ | 3.1816 |
| PC 3 | 0.2600 | $8.0895 \times 10^{-07}$ | 2.3090 |
| PC 4 | 0.3025 | $7.7291 \times 10^{-09}$ | 3.8715 |
| PC 5 | 0.0317 | 0.5539 | 0.7499 |
| PC 6 | 0.0003 | 0.9953 | 1.3037 |
| PC 7 | 0.0796 | 0.1373 | 1.2805 |
| PC 8 | 0.1340 | 0.0121 | 1.1377 |
| PC 9 | 0.0283 | 0.5972 | 1.2028 |
| PC 10 | 0.1876 | 0.0004 | 0.7381 |
| **Chapter (756 Features)** |  |  |  |
| PC 1 | 0.5986 | $2.0784 \times 10^{-35}$ | 4.1839 |
| PC 2 | 0.6914 | $4.6014 \times 10^{-51}$ | 2.9690 |
| PC 3 | 0.3742 | $4.4859 \times 10^{-13}$ | 2.1484 |
| PC 4 | 0.4399 | $5.3695 \times 10^{-18}$ | 3.5000 |
| PC 5 | 0.0494 | 0.3567 | 0.7415 |
| PC 6 | -0.0694 | 0.1955 | 1.3464 |
| PC 7 | 0.0648 | 0.2268 | 1.2850 |
| PC 8 | 0.0897 | 0.0937 | 1.1574 |
| PC 9 | 0.1055 | 0.0485 | 1.1676 |
| PC 10 | 0.1610 | 0.0025 | 0.7481 |

**Table 3:** Top ten feature importances of random forest regression model (Chapter)

| **Feature** | **Importance** |
| --- | --- |
| Black or African American | 0.128 |
| Not Hispanic or Latino | 0.070 |
| Asian | 0.057 |
| Hispanic or Latino | 0.055 |
| Disease of the musculoskeletal system and connective tissue | 0.045 |
| Disease of the nervous system | 0.041 |
| Injury, poisoning and certain other consequences of external causes | 0.041 |
| Certain infectious and parasitic diseases | 0.038 |
| Diseases of the genitourinary system | 0.038 |
| Congenital malformations, deformations and chromosomal abnormalities | 0.036 |
| Mental, Behavioral and Neurodevelopmental disorders | 0.033 |

**Figure 2:**
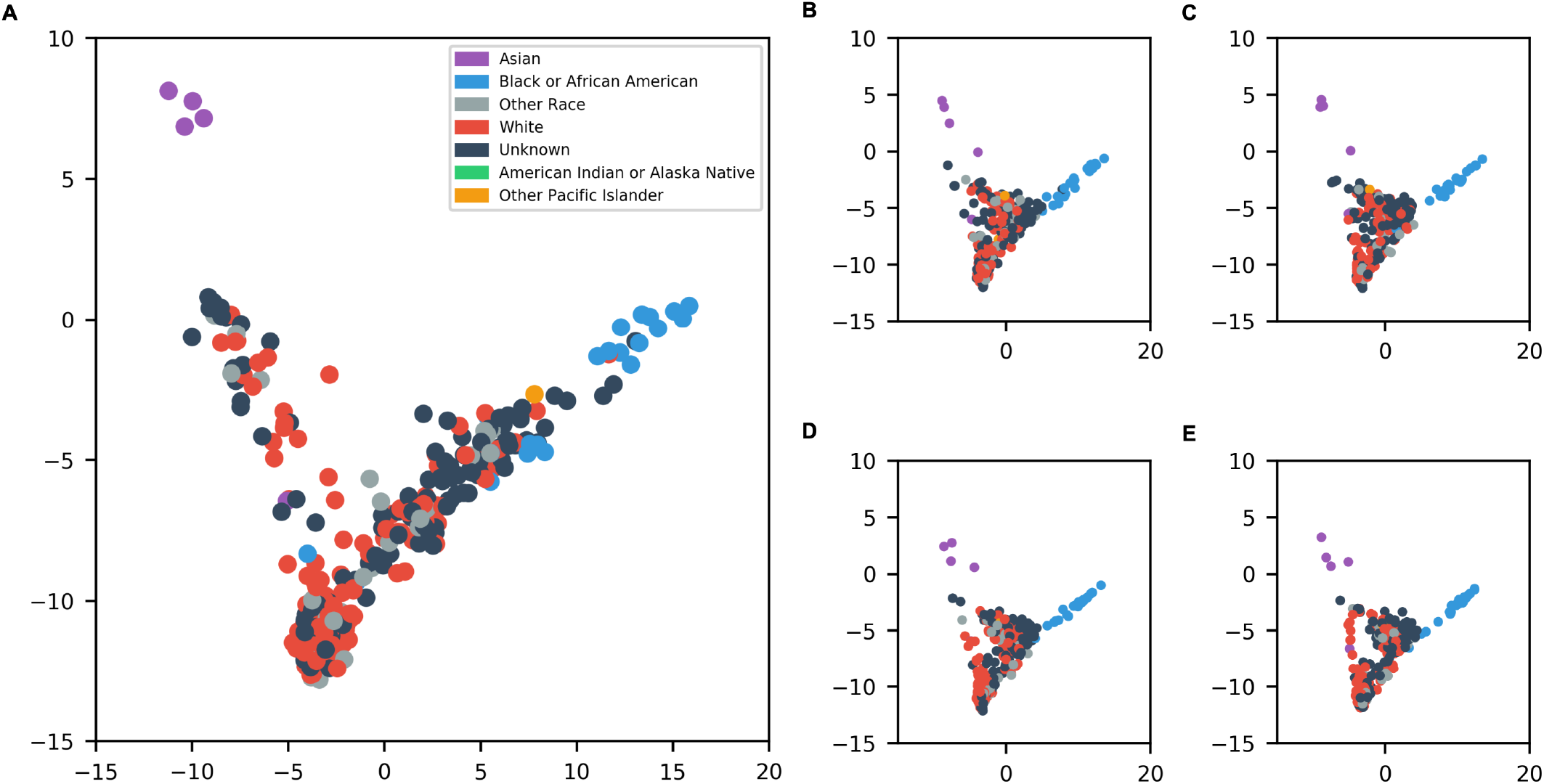
RFR Prediction on the IGM Training Data. Projected PC1 and PC2 (A) and of the RFR predicted PC1 and PC2 using Chapter (B), Block (C), Header (D) and Terminal (D) level features of the IGM training data. Data points are colored according to the patients self reported ancestry.

The RFR algorithm trained on the Chapter level features was evaluated on the validation set of IGM patients (Figure S5). The RFR algorithm predicted the PCs and the projected genetic PCs of the validation set correlated in the same manner as those of the training set data (Figures S5, S6).

### 2.3 Gaussian mixture model predicts population labels

We used a Gaussian mixture (GM) model fit on PCs 1-4 of the 1KGP data from the PCA model and PCs 1-4 of the projection of the HapMap data to assign samples to one of five super populations (EUR, AFR, AMR, EAS, and SAS) (Figure 3). The GM model correctly assigned more than 99% of 1KGP and HapMap data to their super population, each of these regional groups is annotated by their cluster percentage (Table 4).

**Table 4:** Evaluation of Super Population Predictions on 1000 Genomes

| **Super Population** | **Placed Correctly by GMM** |
| --- | --- |
| **African (AFR)** | **99.1%** |
| African Caribbeans in Barbados (ACB) | 100% (96/96) |
| Americans of African Ancestry in SW USA (ASW) | 94.2% (97/103) |
| Esan in Nigeria (ESN) | 100% (99/99) |
| Gambian in Western Divisions in the Gambia (GWD) | 100% (113/113) |
| Luhya in Webuye, Kenya (LWK) | 100% (121/121) |
| Maasai in Kinyawa, Kenya (MKK) | 100% (184/184) |
| Mende in Sierra Leone (MSL) | 100% (85/85) |
| Yoruba in Ibadan, Nigeria (YRI) | 98.6% (209/212) |
| **Admixed American (AMR)** | **97.4%** |
| Colombians from Medellin, Colombia (CLM) | 96.8% (91/94) |
| Mexican Ancestry from Los Angeles USA (MXL) | 96.9% (94/97) |
| Peruvians from Lima, Peru (PEL) | 100% (85/85) |
| Puerto Ricans from Puerto Rico (PUR) | 96.2% (100/104) |
| **East Asian (EAS)** | **99.9%** |
| Chinese Dai in Xishuangbanna, China (CDX) | 100% (93/93) |
| Han Chinese in Beijing, China (CHB) | 99.3% (146/147) |
| Chinese in Metropolitan Denver, Colorado (CHD) | 100% (109/109) |
| Southern Han Chinese (CHS) | 100% (105/105) |
| Japanese in Tokyo, Japan (JPT) | 100% (120/120) |
| Kinh in Ho Chi Minh City, Vietnam (KHV) | 100% (99/99) |
| **European (EUR)** | **98.3%** |
| Utah Residents (CEPH) with Northern and Western European Ancestry (CEU) | 97.2% (174/179) |
| British in England and Scotland (GBR) | 100% (91/91) |
| Finnish in Finland (FIN) | 100% (99/99) |
| Iberian Population in Spain (IBS) | 96.3% (103/107) |
| Toscani in Italia (TSI) | 99.1% (112/113) |
| **South Asian (SAS)** | **100% (85/85)** |
| Bengali from Bangladesh (BEB) | 100% (86/86) |
| Gujarati Indian from Houston, Texas (GIH) | 100% (113/113) |
| Indian Telugu from the UK (ITU) | 100% (102/102) |
| Punjabi from Lahore, Pakistan (PJL) | 100% (96/96) |
| Sri Lankan Tamil from the UK (STU) | 100% (102/102) |

**Figure 3:**
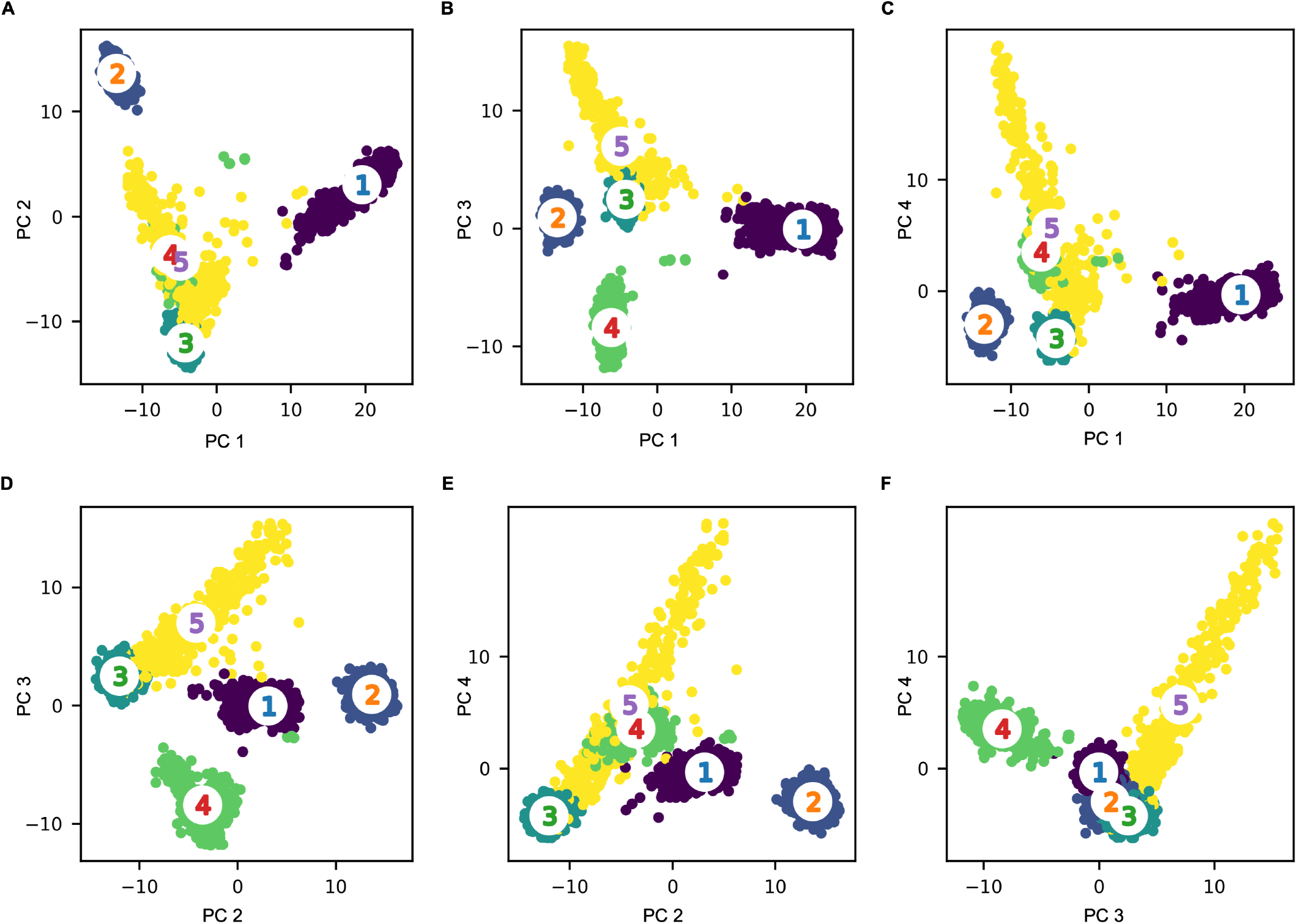
GM Model of 1KGP and HapMap Data. PC1 and PC2 (A), PC1 and PC3 (B), PC1 and PC4 (C), PC2 and PC3 (D), PC2 and PC4 (E), and PC3 and PC4 (F) of the 1KGP and independent HapMap data colored according to assigned GM model cluster.

### 2.4 Ancestry prediction assign labels to ambiguous patient populations

Ancestry information incomplete in the EHR, more than 75% of the patients in the CU/NYIMC EHR have unknown race and more than 85% have unknown ethnicity (Table 1). Using our pipeline (the RFR algorithm and the GM model), we are able to assign an ancestry label to all New York City patients (N=2,673,627) (Table 5). New York City patients that reported unknown or other are overwhelmingly classified as Ad Mixed American based on this pipeline.

**Table 5:** Predicted Super Population and Self-Reported Race and Ethnicity in NYC patients

|  | **African**<br>**(AFR)** | **Ad Mixed**<br>**American**<br>**(AMR)** | **East Asian**<br>**(EAS)** | **European**<br>**(EUR)** | **South**<br>**Asian**<br>**(SAS)** |
| --- | --- | --- | --- | --- | --- |
| **Total** | 197,240 | 2,264,514 | 22,565 | 187,230 | 1078 |
| **Self Reported Race** |  |  |  |  |  |
| American Indian or Native American | 3.81% | **93.05%** | 0.27% | 2.87% | 0% |
| Asian | 0.23% | 3.04% | **92.07%** | 0.25% | 4.40% |
| Black | **87.28%** | 12.50% | 0.01% | 0.20 | <0.01% |
| Pacific Islander | 8.65% | **82.25%** | 0.95% | 7.88 | 0.27% |
| White | 0.03% | **69.25%** | <0.01% | **30.72%** | 0% |
| Unknown | 0.02% | **99.98%** | <0.01% | <0.01% | <0.01% |
| Other Race | <0.01% | **99.58%** | <0.01% | 0.42% | 0% |
| **Self Reported Ethnicity** |  |  |  |  |  |
| Hispanic | 0.83% | **99.17%** | 0% | 0 | 0% |
| Non-hispanic | 8.73% | **81.75%** | 1.02% | 8.45 | 0.05 |

### 2.5 Ancestry predictions reflect demographics by zip code

In order to analyze the power of our pipeline, we compared the distribution of Black or African American (AFR), East Asian (EAS) and EUR (European) predicted super population across each zip code in New York City with the most recent demographic data (Figure 4). The demographic distribution by zip code strongly correlates with the US Census Estimates^13^ (R2 = 0.946, p=1.45*×*10^*−*93^ (AFR and Black), R2=0.900, p=1.97*×*10^*−*69^ (EAS and Asian), R2 = 0.839, p=2.64*×*10^*−*51^ (EUR and White)). Our pipeline identified regions with a large fraction of AFR ancestry, such as 10460, 10469, 10475, 10039, and 11434, EAS ancestry, such as 11354, 11355, 10002, and 11220, and with EUR ancestry, such as 11202, 11693, 10471, 10463, and 10282.

**Figure 4:**
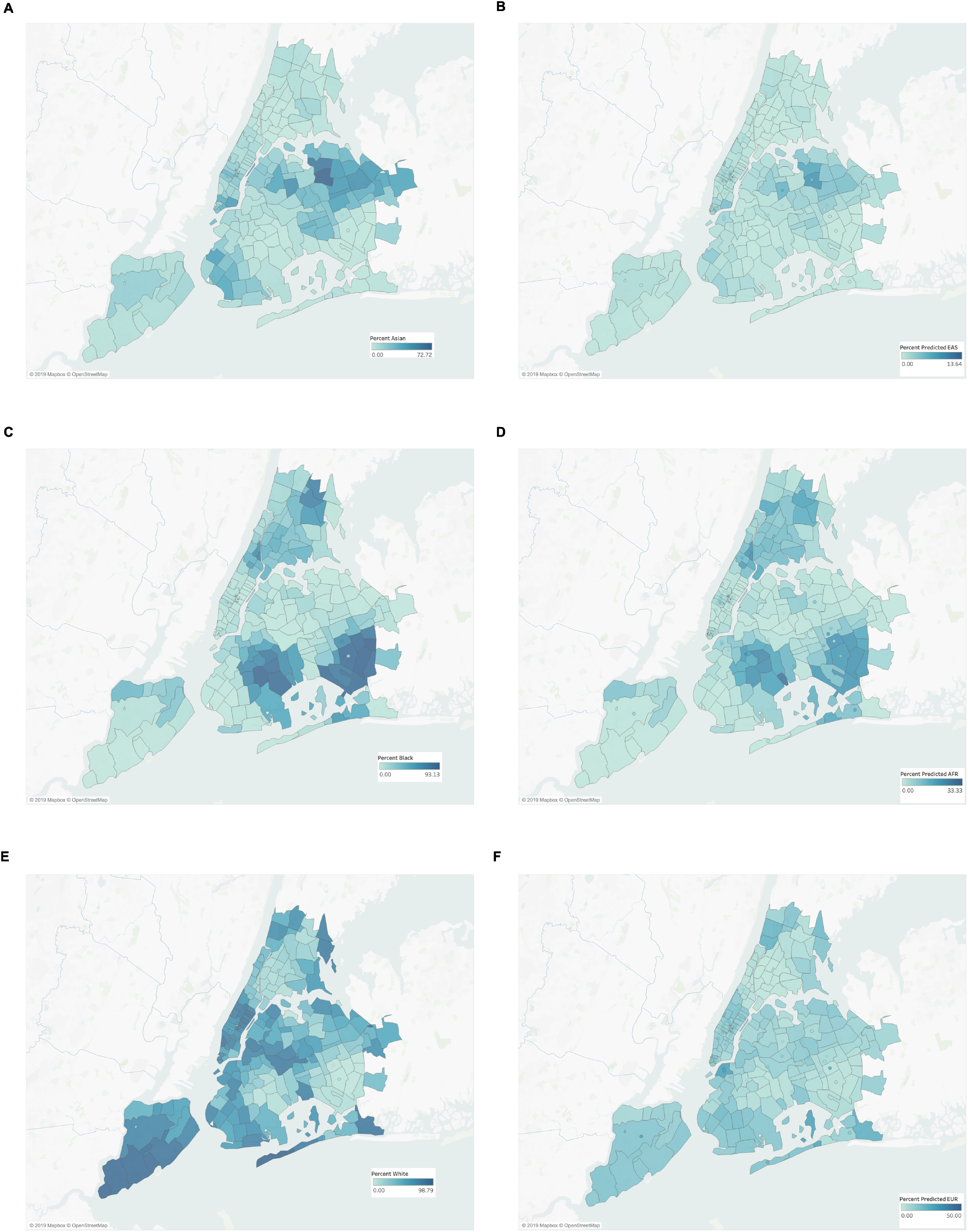
Ancestry of New York City Residents and CU/NYPIMC Patients by Zip Code. Percent of NYC residents who identified as Asian (A), Black (C), and White (E) according to recent US Census Estimation and CU/NYPIMC who, according to our pipeline, are predicted to be of East Asian (B), Black or African American (D), and European (F) ancestry by zip code. Only patients with a NYC zip code are included in these maps.

### 2.6 Using ancestry predictions to analyze disease incidence rates

To further analyze the power of our pipeline, we compared the incidence rate of different diseases by predicted super population (Table 6, S2). Based on the patient population, attention deficit and hyperactivity disorder (ADHD), Huntington’s disease, obesity, and uncontrolled type 2 diabetes is more prevalent in AFR or EUR ancestry populations compared to the EAS ancestry population. Polydactyly and sickle cell trait is more prevalent in the AFR ancestry population compared to EAS or EUR ancestry populations. Celiac disease and cystic fibrosis is more prevalent in the EUR ancestry population compared to AFR or EAS ancestry populations. Based on the patient population, trisomy 13 (Patau syndrome), trisomy 18 (Edwards’ syndrome) and trisomy 21 (Down syndrome) is equally prevalent in AFR, EAS, and EUR ancestry populations.

**Table 6:**
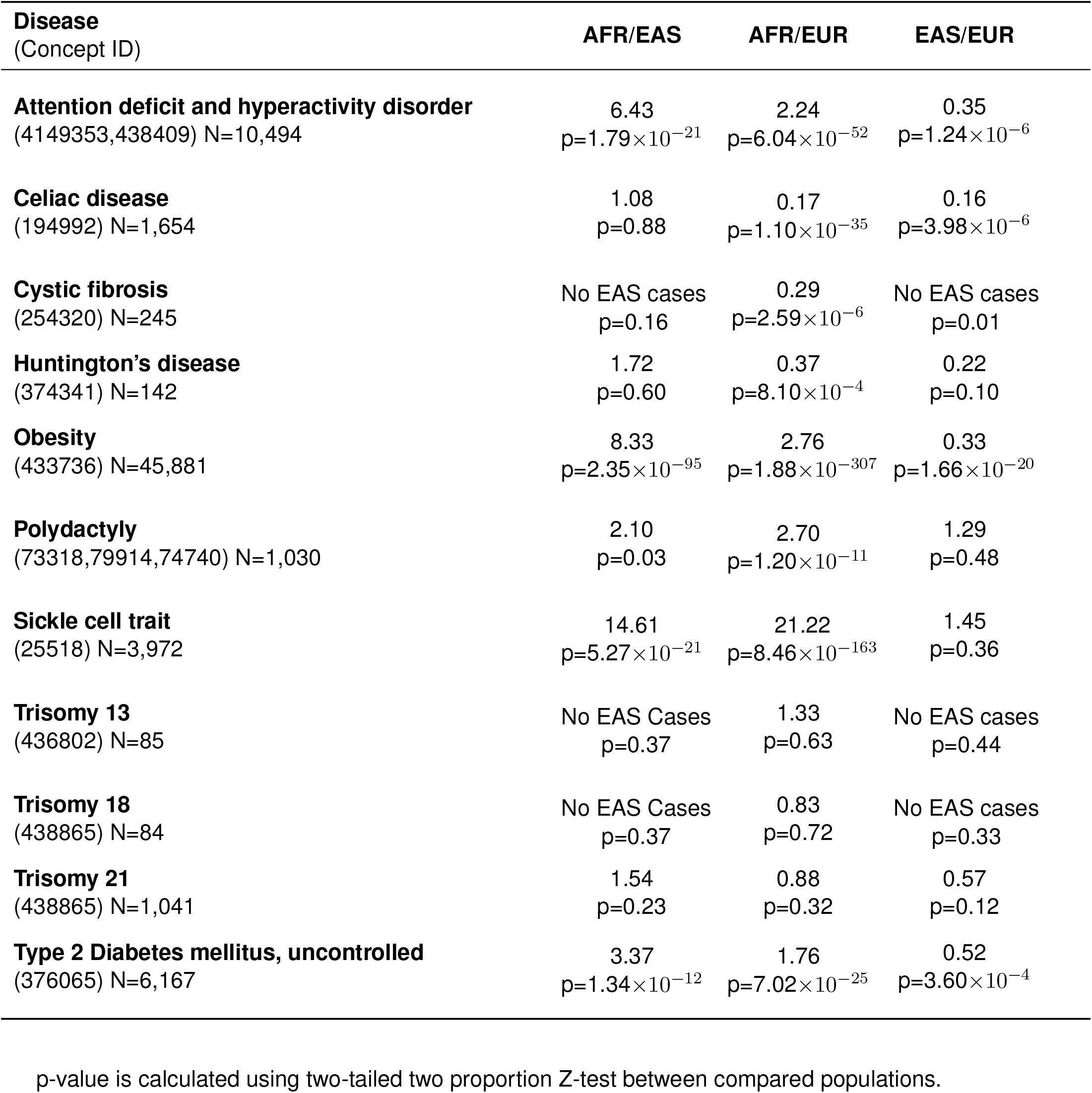
Relative Prevalence Rates of Diseases by Predicted Ancestry

| **Disease**<br>(Concept ID) | **AFR/EAS** | **AFR/EUR** | **EAS/EUR** |
| --- | --- | --- | --- |
| **Attention deficit and hyperactivity disorder**<br>(4149353,438409) N=10,494 | 6.43<br>$p=1.79 \times 10^{-21}$ | 2.24<br>$p=6.04 \times 10^{-52}$ | 0.35<br>$p=1.24 \times 10^{-6}$ |
| **Celiac disease**<br>(194992) N=1,654 | 1.08<br>$p=0.88$ | 0.17<br>$p=1.10 \times 10^{-35}$ | 0.16<br>$p=3.98 \times 10^{-6}$ |
| **Cystic fibrosis**<br>(254320) N=245 | No EAS cases<br>$p=0.16$ | 0.29<br>$p=2.59 \times 10^{-6}$ | No EAS cases<br>$p=0.01$ |
| **Huntington's disease**<br>(374341) N=142 | 1.72<br>$p=0.60$ | 0.37<br>$p=8.10 \times 10^{-4}$ | 0.22<br>$p=0.10$ |
| **Obesity**<br>(433736) N=45,881 | 8.33<br>$p=2.35 \times 10^{-95}$ | 2.76<br>$p=1.88 \times 10^{-307}$ | 0.33<br>$p=1.66 \times 10^{-20}$ |
| **Polydactyly**<br>(73318,79914,74740) N=1,030 | 2.10<br>$p=0.03$ | 2.70<br>$p=1.20 \times 10^{-11}$ | 1.29<br>$p=0.48$ |
| **Sickle cell trait**<br>(25518) N=3,972 | 14.61<br>$p=5.27 \times 10^{-21}$ | 21.22<br>$p=8.46 \times 10^{-163}$ | 1.45<br>$p=0.36$ |
| **Trisomy 13**<br>(436802) N=85 | No EAS Cases<br>$p=0.37$ | 1.33<br>$p=0.63$ | No EAS cases<br>$p=0.44$ |
| **Trisomy 18**<br>(438865) N=84 | No EAS Cases<br>$p=0.37$ | 0.83<br>$p=0.72$ | No EAS cases<br>$p=0.33$ |
| **Trisomy 21**<br>(438865) N=1,041 | 1.54<br>$p=0.23$ | 0.88<br>$p=0.32$ | 0.57<br>$p=0.12$ |
| **Type 2 Diabetes mellitus, uncontrolled**<br>(376065) N=6,167 | 3.37<br>$p=1.34 \times 10^{-12}$ | 1.76<br>$p=7.02 \times 10^{-25}$ | 0.52<br>$p=3.60 \times 10^{-4}$ |

## 3 Discussion

The advances in genetics, both in research and the clinic, are significant and continuously expanding. Newly available clinical data present an equally transformative opportunity. These data however, are limited by issues of missingness and inaccuracy^14^. If properly addressed, the EHR could be used as a resource to identify disease and treatment subgroups that inform recruitment criteria for genetic studies and to support genetic research^15^. Currently, these data are not being used to their full capacity, mainly due to the scarcity of linked genetic data. Here, we analyzed data from the EHR in conjunction with existing genetic data at CU/NYPIMC that are available for a small subset of our patients. We then utilized common data elements captured by the EHR to predict patient ancestry for over 2.6 million patients, who reside in New York City.

In this study, we collected anonymized genetic data from individuals that consented to have their results used for research and merged these datasets, using 5,758 common variants from the 1KGP populations^16^ as a reference. We derived a PCA model based on the 1KGP, and applied the model to the IGM genetic data and matched the genetic data to clinical information from the EHR for the same individuals. The PCA model was limited by a small number of samples with genetic data (2,504 samples from 1KGP, 650 samples from HapMap, and 350 samples from IGM) and a limited set of common variants. Future work may incorporate larger genetic datasets such as the Exome Aggregation Consortium (ExAC; 60,706 unrelated samples with 7,404,909 high-quality variants)^17^, or the more recent genome Aggregation Database (gnomAD; 123,136 exome sequences and 15,496 whole-genome sequences), to include more rare ancestry informing markers. We then trained a machine learning algorithm (random forest regression) to predict PC values using the clinical data. While this model generally yielded accurate predictions, we found that the model had more difficulty correctly predicting the extremes of the PC space (e.g. patients that reported Asian ancestry). Accuracy of predictions are also affected by demographic differences of patients used for training the machine learning algorithm versus the population of the EHR. In the future, we will attempt to address these limitations by expanding the population of patients with linked genetic data and by using more adaptable models^18^.

We used a GM Model to assign ancestry labels to the predicted PC values for the EHR population. These ancestry labels are especially informative for patients for whom rave and ethnicity data is not available or collected. Furthermore, the options that are available for self-reporting are often insufficient. This is also important for researchers using the EHR for data mining of populations believed to share genetic factors responsible for a disease or drug response phenotype. We were able to assign ancestry information to 1,869,110 New York City patients with unknown self reported race or that report other. In addition to ancestry population classification, each of the 2.6 million patients also have PC values for PCs 1-4 predicted and available for further interpretation. While the labels are important for validation and visualization, but the PC values will be more useful than categories. Proximity in PC space may cross ancestry labels, but still indicative of genetic similarity relevant for disease susceptibility and drug sensitivity.

We used the assigned ancestry to analyze the demographics of New York City by zip code. The population distribution using the predicted ancestry from our pipeline by zip code highly correlated with the ancestry distribution from the US Census Estimates^13^. We further identified disease incidence rates between the predicted ancestry groups. The incidence rate in the patients at CU/NYPIMC by ancestry group recapitulates the observed prevalence between ancestry subgroups for Celiac disease^19^, cystic fibrosis^20^, Huntington’s disease^21^, obesity^22^, polydactyly^23^, sickle cell trait^24^, and type 2 diabetes^25^. The observed incidence rate in the New York City patients for attention deficit and hyperactivity disorder does not agree with the past observations^26^, which documented a lower prevalence in African American and Hispanic children compared to white children.

The incidence rates of trisomy 13 and 18 in the patients at CU/NYPIMC is not significantly different between AFR, EAS, and EUR ancestry groups agreeing with previous analysis^27,28^, which adds power to the applicability of this pipeline. The incidence rates of trisomy 21 in the patients at CU/NYPIMC is not significantly different between AFR, EAS, and EUR ancestry groups, which contrasts with previous observations identifying a higher prevalence in African Americans compared to Asian and white populations^28^.

## 4 Conclusion

We have developed a new pipeline of two machine learning algorithms to predict genetic ancestry using only clinical data. These predictions can be used to assign ancestry labels to patients in the electronic health record for whom genetic data was previously unavailable.

This information is helpful for identifying subgroups that are likely to be ancestry specific.

## 5 Materials and Methods

### 5.1 Acquiring genetic data at Columbia

We obtained WES data for patients that consented to have their results used for research from the Institute for Genomic Medicine at CUIMC/NYPIMC.

All datasets were processed using PLINK^29^ and custom Python scripts. All SNPs were filtered to exclude indels and be observed at a minor allele frequency of at least 10% and a genotyping rate of at least 90%, and were normalized to a set of shared variants.

### 5.2 Fitting principal components analysis (PCA) model

To project our samples into PC space defined by 1KGP, we first identified all shared common SNVs between our datasets, the 1KGP, and HapMap (excluding redundant samples between the latter two datasets)^30^. We began by converting the HapMap variants to the University of California Santa Cruz (UCSC) 2009 Genome Reference Consortium human reference sequence build (GRCh37/hg19) using the liftOver tool^31^, enabling all samples to be compared. We isolated the shared SNV among this group and our sequencing data. We then fit a PCA model to 1KGP data using the shared SNVs and then applied this model to HapMap and the IGM training set. Only the 1KGP data was used to train the PCA model. All code is available in a publicly accessible git repository.

### 5.3 Matching to clinical population

We used a common patient identifier to determine the subset of patients with genetic data who also had available clinical data (demographics and conditions) in our CDW. Of the 726 samples available, the set of samples available before May 2017 was used as the training set for the RFR algorithm (N=350) and the remainder of the samples were used as the validation set for the RFR algorithm (N=376). The study and use of all clinical and genetic data was approved by the Columbia Institutional Review Board. See Table 1 for a demographic description of the cohort.

### 5.4 Training random forest model to predict principal components

We trained a random forest regression model to predict PC values from clinical data using the IGM training set. We extracted diagnosis billing codes for each patient to use as features for machine learning. These codes represent what the patient was billed for for their visit to the hospital and provides a general overview of their state. These data are coded in either the ICD-9 (before November 2015) or ICD-10 vocabularies (after November 2015). We mapped ICD-9 codes in the EHR to ICD-10 using the General Equivalence Mappings. We used the heirarchical structure of the ICD-10 codes to reduce the number of features that the model was trained on. We defined features at four levels of the ICD-10 hierarchy (Terminal, Header, Block, Chapter). These levels are in order of decreasing specificity; for example: *Terminal*: (J06.0) Acute laryngopharyngitis; *Header*: (J06) Acute upper respiratory infections of multiple and unspecified sites; *Block*: (J00-J06) Acute upper respiratory infections; *Chapter*: (J00-J99) Diseases of the respiratory system. We used these conditions and patient demographics (self-reported race and ethnicity) as features to predict the first ten principal components for each patient in the training set. We trained the model using 735 demographic features plus the conditions from each of the four levels in the hierarchy.

### 5.5 Evaluating and optimizing model performance

We evaluated the random forest regression model performance using out-of-bag (OOB) predictions. OOB produces robust estimates of performance on par to cross validation^32^. For each principal component, we calculated the Pearson correlation and RMSE between the actual and predicted PCs.

### 5.6 Predicting population labels

We used a GM model to predict ancestry labels for each individual in the EHR using the first four PC values. We evaluated the predicted labels using the known labels for the individuals in 1KGP. All datasets were processed using custom Python scripts and are available in a public git repository.

### 5.7 Mapping Population Demographics

To better visualize our demographics, we mapped individuals with New York City addresses on a map using their predicted ancestry labels assigned with our pipeline, and compared this to the demographic information collected through the US Census^13^. Zip code alignment and visualization of maps was done using Tableau^33^.

### 5.8 Analyzing Disease Incidence Rates

To analyze our pipeline, we identified individuals with New York City address and a concept id for the particular disease of interest cross referencing for the GM model assigned ancestry. For diseases with multiple concept ids, individuals with any of the concept ids were counted and filtered to preventing inaccurate counting.

## 6 Supplementary Material

**Table S1:** Random forest regression model performance with feature levels

| Features | Pearson correlation | p-value | RMSE |
| --- | --- | --- | --- |
| **Demographics** |  |  |  |
| PC 1 | 0.5531 | $1.9304 \times 10^{-29}$ | 4.3404 |
| PC 2 | 0.6348 | $7.1646 \times 10^{-41}$ | 3.1741 |
| PC 3 | 0.3637 | $2.1961 \times 10^{-12}$ | 2.1001 |
| PC 4 | 0.4009 | $6.0363 \times 10^{-15}$ | 3.4773 |
| **Conditions** |  |  |  |
| PC 1 | 0.3087 | $3.6830 \times 10^{-09}$ | 5.1266 |
| PC 2 | 0.2839 | $6.5534 \times 10^{-08}$ | 4.1728 |
| PC 3 | 0.0568 | 0.2894 | 2.3891 |
| PC 4 | 0.2372 | $7.2686 \times 10^{-06}$ | 3.8227 |
| **Demographics and Conditions** |  |  |  |
| PC 1 | 0.5986 | $2.0784 \times 10^{-35}$ | 4.1839 |
| PC 2 | 0.6914 | $4.6014 \times 10^{-51}$ | 2.9690 |
| PC 3 | 0.3742 | $4.4859 \times 10^{-13}$ | 2.1484 |
| PC 4 | 0.4399 | $5.3695 \times 10^{-18}$ | 3.5000 |

**Table S2:** Disease incidences per 10,000 individuals for AFR, EAS, and EUR Ancestry (± standard error)

| **Disease**<br>Concept ID | **AFR**<br>N=197,240 | **EAS**<br>N=22,565 | **EUR**<br>N=187,230 |
| --- | --- | --- | --- |
| **Attention deficit and hyperactivity disorder**<br>(4149353,438409) | 57.0±0.5 | 8.9±0.2 | 25.4±0.3 |
| **Celiac disease**<br>(194992) | 2.4±0.1 | 2.2±0.1 | 13.7±0.2 |
| **Cystic fibrosis**<br>(254320) | 0.9±0.1 | 0 | 2.9±0.1 |
| **Huntington disease**<br>(374341) | 0.8±0.1 | 0.4±0.04 | 2.0±0.1 |
| **Obesity**<br>(433736) | 243.6±0.9 | 29.2±0.3 | 88.1±0.6 |
| **Sickle cell trait**<br>(25518) | 45.3±0.4 | 3.1±0.2 | 2.1±0.1 |
| **Polydactyly**<br>(73318,79914,74740) | 8.4±0.2 | 4.0±0.1 | 3.1±0.1 |
| **Trisomy 13**<br>(436802) | 0.4±0.04 | 0 | 0.3±0.03 |
| **Trisomy 18**<br>(438865) | 0.4±0.04 | 0 | 0.4±0.04 |
| **Trisomy 21**<br>(439125) | 5.5±0.1 | 3.5±0.1 | 6.2±0.2 |
| **Type 2 Diabetes mellitus, uncontrolled**<br>(376065) | 46.3±0.4 | 13.7±0.2 | 26.2±0.3 |

**Figure S1:**
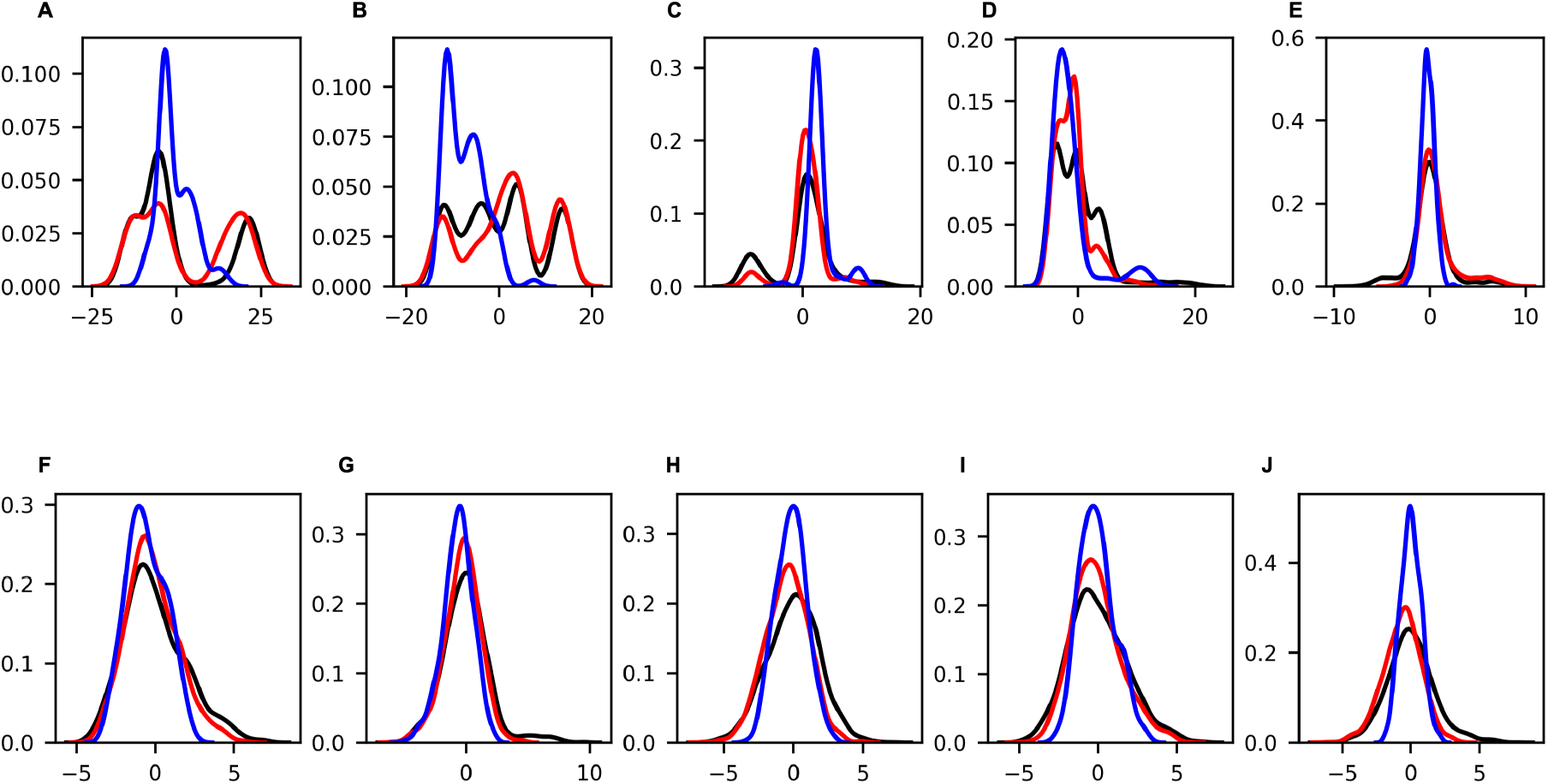
Distribution of PCA data. Density plots of the distribution of values for each PC (A-J: PC1-10, respectively) for the 1KGP data (black), the HapMap projection (red), and the IGM training set projection (blue).

**Figure S2:**
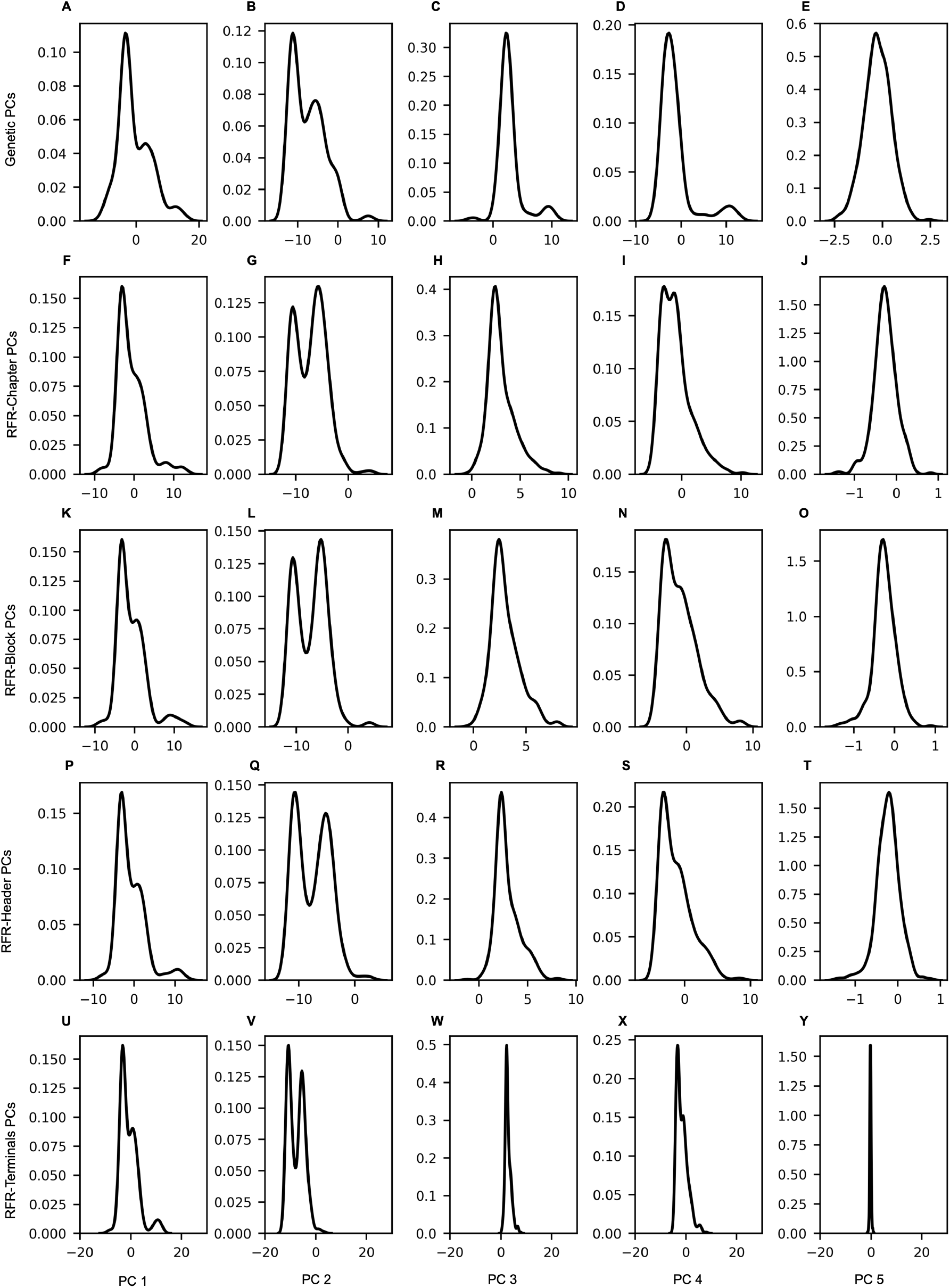
Distribution of PCs for IGM Training Set. Density plots of the distribution of values for the projected PCs (A-E: PC1-5, respectively) and the for the predicted PCs by the RFR algorithm using Chapter (F-J: PC1-5, respectively), Block (K-O: PC1-5, respectively), Header(P-T: PC1-5, respectively), and Terminal (U-Y: PC1-5, respectively) level features.

**Figure S3:**
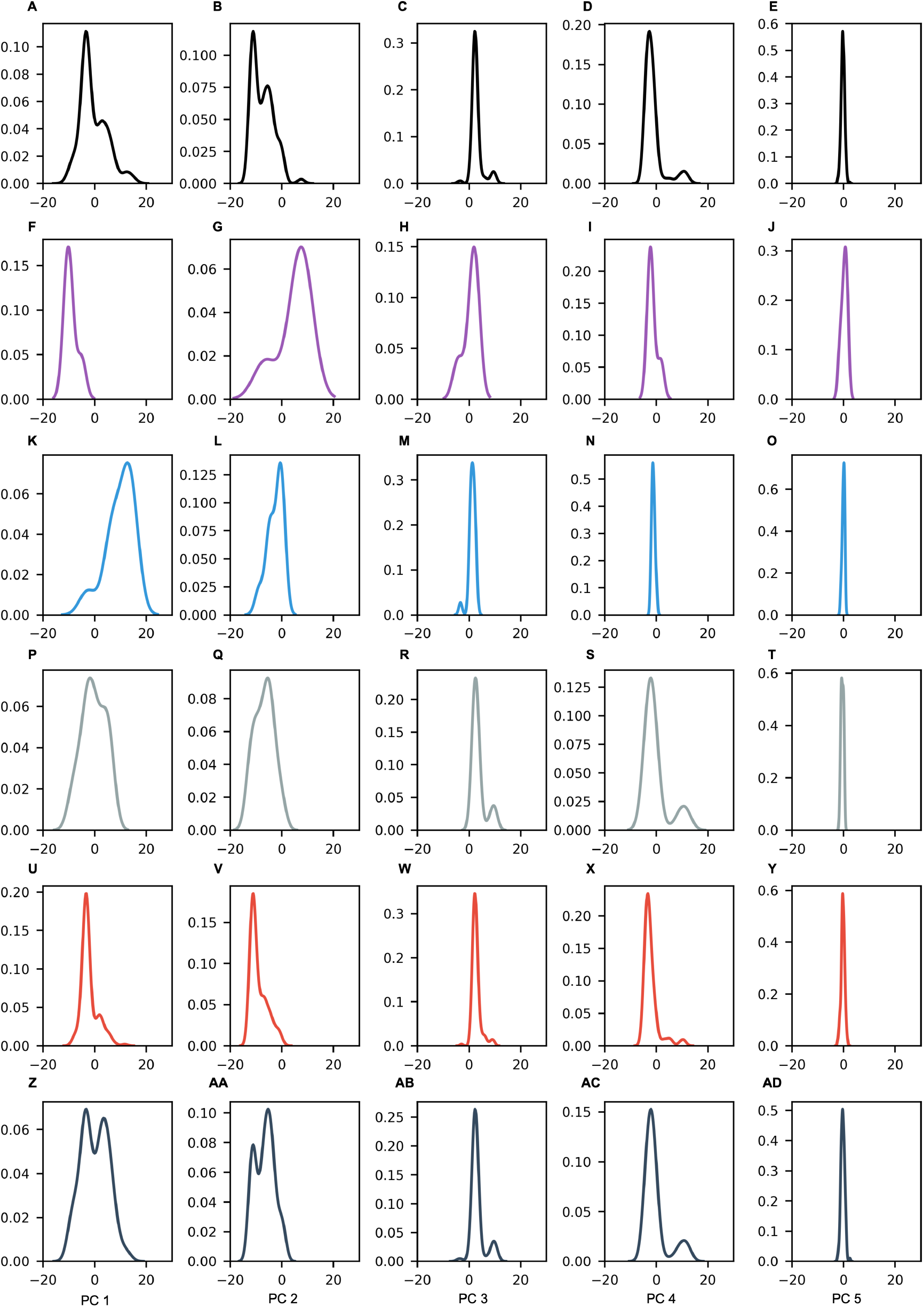
Distribution of Projected PCs by Self Reported Race. Density plots of the distribution of values for the projected PCs for all patients in the training set (A-E: PC1-5, respectively), and for patients who self reported Asian (F-J: PC1-5, respectively), Black (K-O: PC1-5, respectively), Other (P-T: PC1-5, respectively), White (U-Y: PC1-5, respectively), and Unknown (Z-AD: PC1-5, respectively) ancestry.

**Figure S4:**
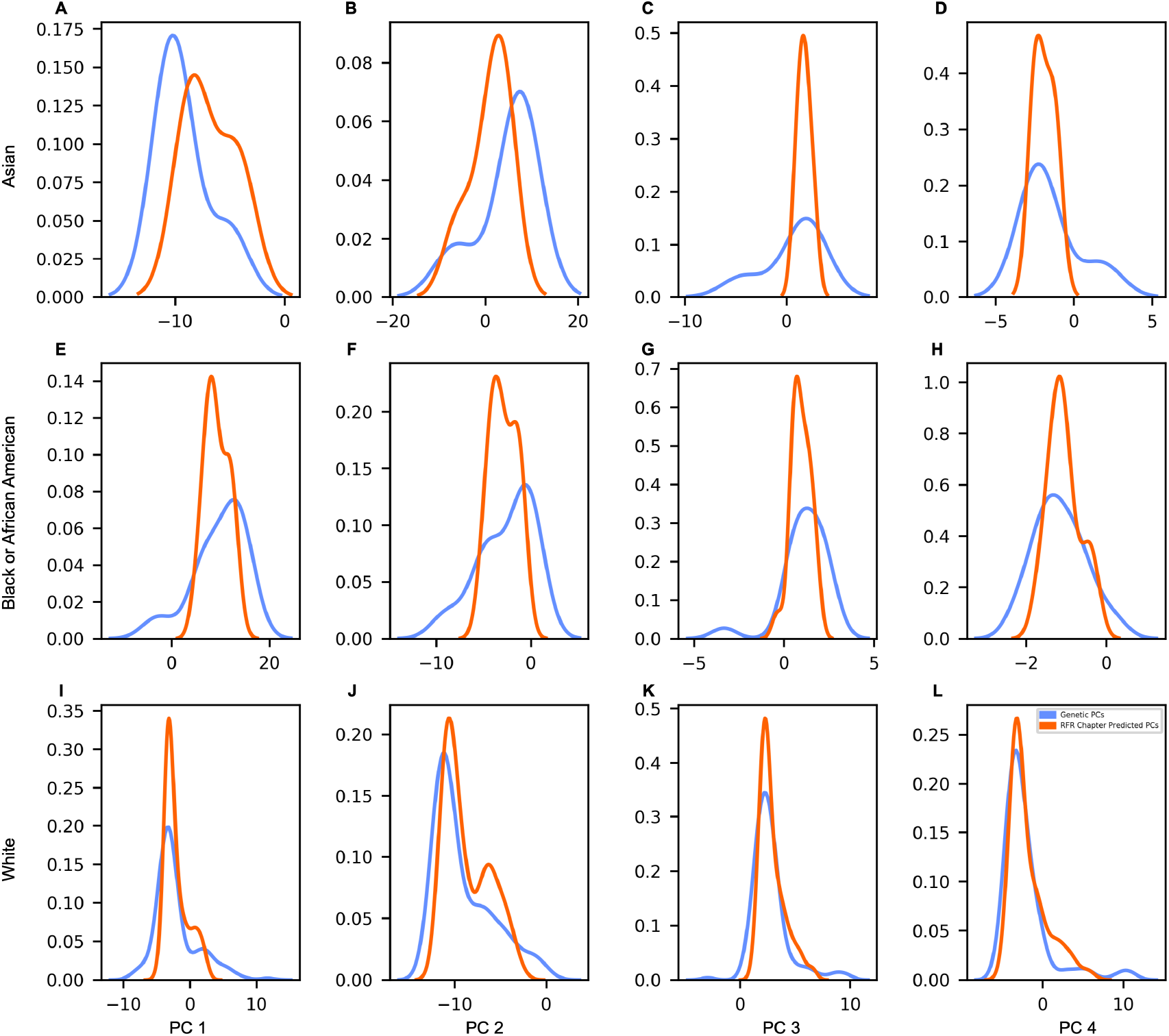
Distribution of Projected and Predicted PCs by Self Reported Race. Density plots of the distribution of values for the projected PCs (blue) and PCs predicted by the RFR algorithm trained on Chapter level features (orange) for IGM training set patients who self reported as Asian (A-D: PC1-4, respectively), Black (E-H: PC1-4, respectively), and White (I-L: PC1-4) ancestry.

**Figure S5:**
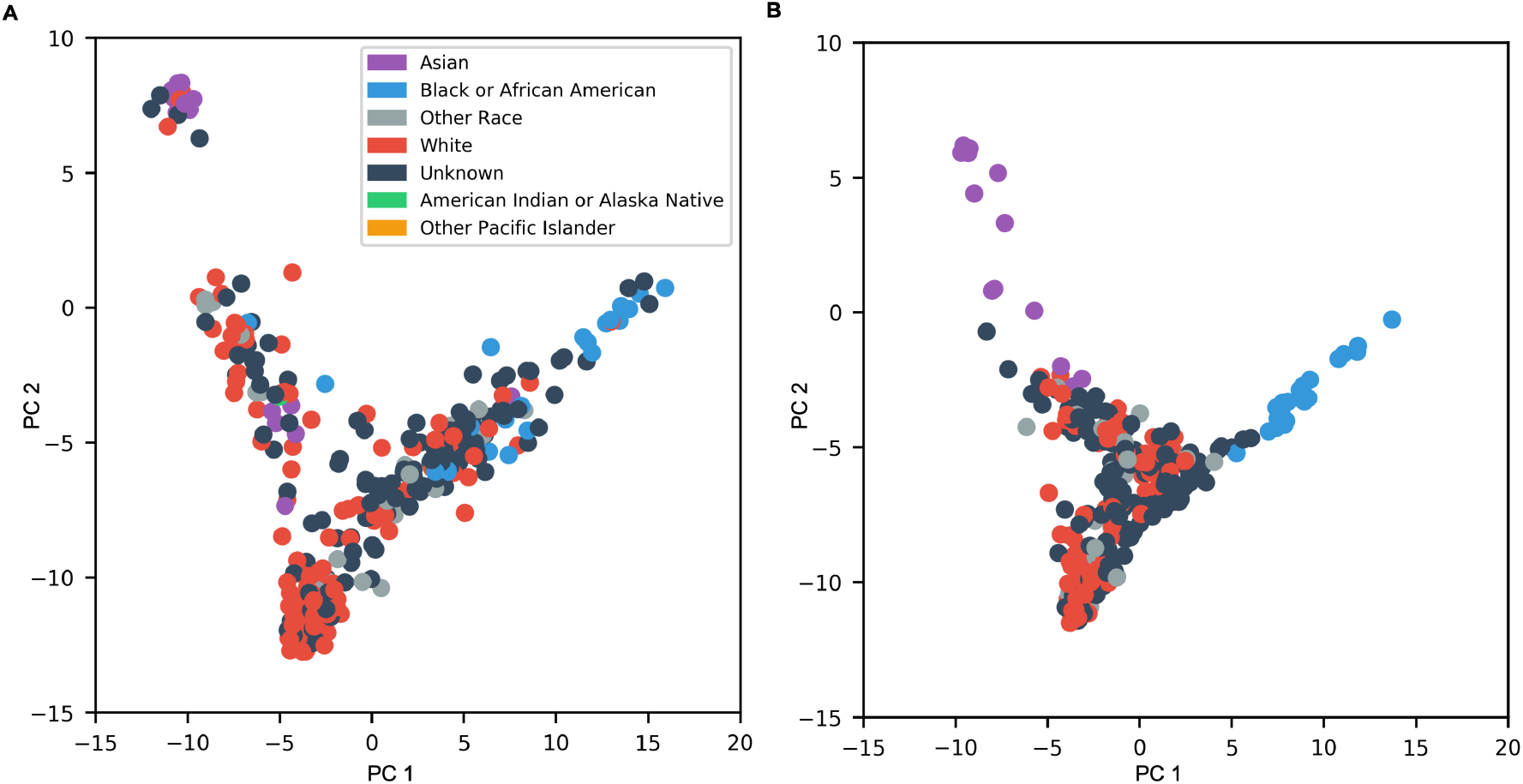
RFR Prediction on the IGM Validation Set. Projected PC1 and PC2 (A) and the RFR predicted PC1 and PC2 using Chapter level features (B) for the IGM validation set. Data points are colored according to the patients self reported ancestry.

**Figure S6:**
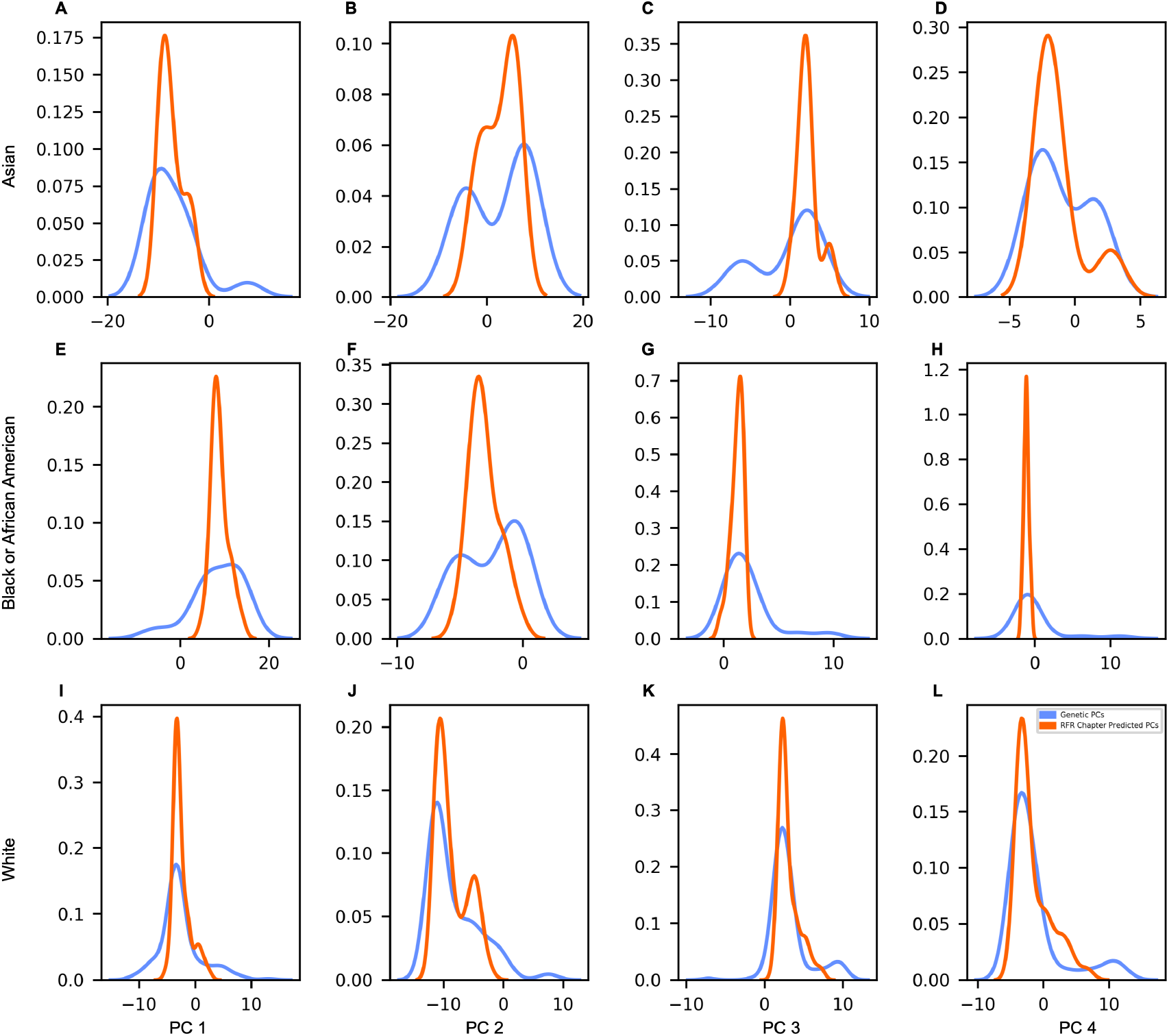
Distribution of Projected and Predicted PCs by Self Reported Race. Density plots of the distribution of values for the projected PCs (blue) and PCs predicted by the RFR algorithm trained on Chapter level features (orange) for IGM validation set patients who self reported as Asian (A-D: PC1-4, respectively), Black (E-H: PC1-4, respectively), and White (I-L: PC1-4) ancestry.

**Figure S7:**
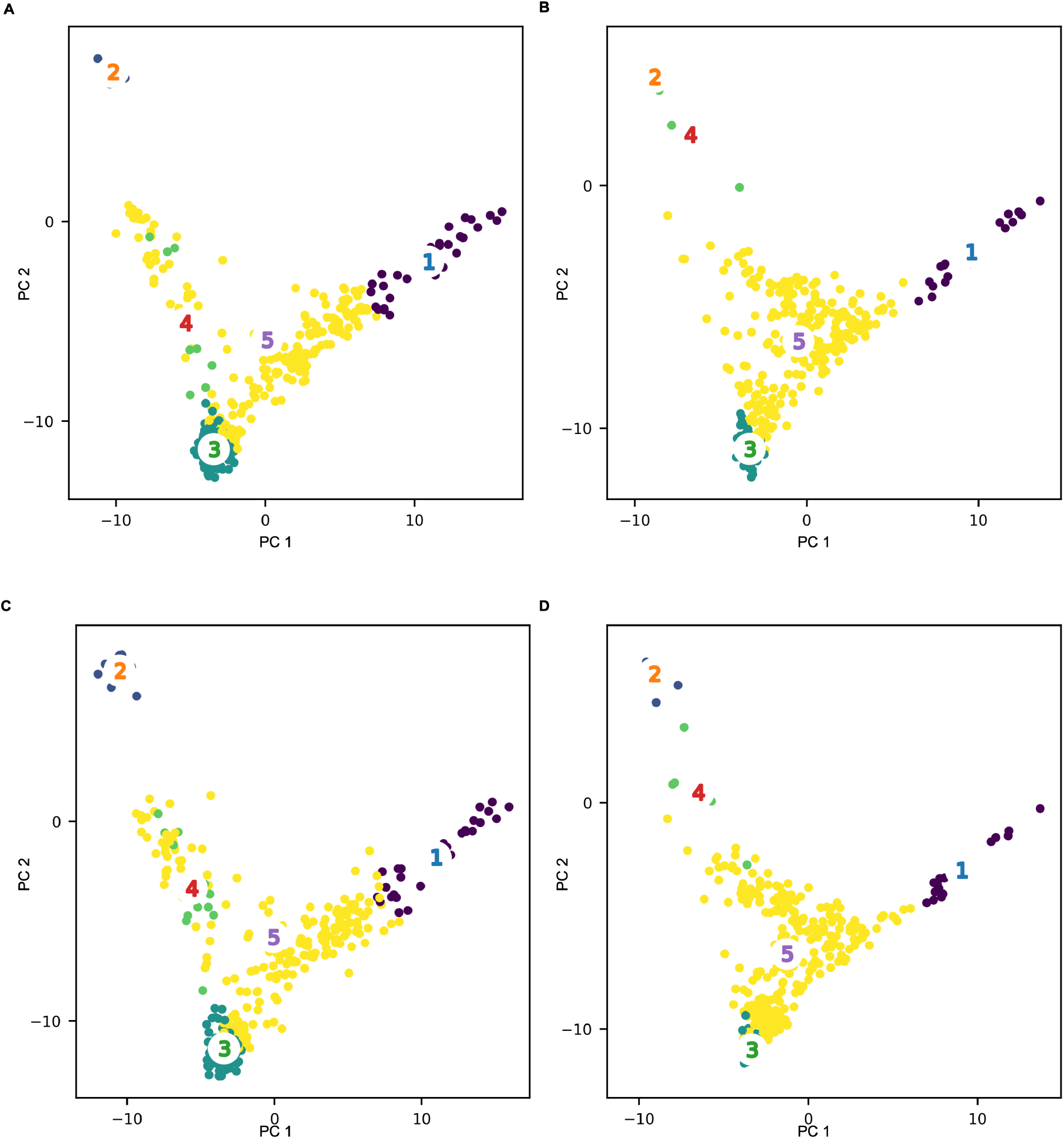
GM Model Applied to the IGM Data. GM model applied to the projected (A, C) and predicted (B, D) PCs for the IGM training set (A, B) and and validation set (C,D) colored according to assigned GM model cluster.

